# A single nucleotide change in the *polC* DNA polymerase III in *Clostridium thermocellum* is sufficient to create a hypermutator phenotype

**DOI:** 10.1101/2021.07.30.454558

**Authors:** Anthony Lanahan, Kamila Zakowicz, Liang Tian, Daniel G. Olson, Lee R. Lynd

## Abstract

*Clostridium thermocellum* is a thermophilic, anaerobic, bacterium that natively ferments cellulose to ethanol, and is a candidate for cellulosic biofuel production. Recently, we identified a hypermutator strain of *C. thermocellum* with a C669Y mutation in the *polC* gene, which encodes a DNA polymerase III enzyme. Here we reintroduce this mutation using recently-developed CRISPR tools to demonstrate that this mutation is sufficient to recreate the hypermutator phenotype. The resulting strain shows an approximately 30-fold increase in the mutation rate. This mutation appears to function by interfering with metal ion coordination in the PHP domain responsible for proofreading. The ability to selectively increase the mutation rate in *C. thermocellum* is a useful tool for future directed evolution experiments.

**Importance:** Cellulosic biofuels are a promising approach to decarbonize the heavy duty transportation sector. A longstanding barrier to cost-effective cellulosic biofuel production is the recalcitrance of the material to solubilization. Native cellulose-consuming organisms, such as *Clostridium thermocellum*, are promising candidates for cellulosic biofuel production, however they often need to be genetically modified to improve product formation. One approach is adaptive laboratory evolution. Our findings demonstrate a way to increase the mutation rate in this industrially-relevant organism, which can reduce the time needed for adaptive evolution experiments.

## Introduction

*Clostridium thermocellum* (aka *Acetivibrio thermocellus, Hungateiclostridium thermocellum,* and *Ruminiclostridium thermocellum*) is a thermophilic, anaerobic bacterium that can ferment crystalline cellulose to ethanol and has attracted interest as a candidate for cellulosic biofuel production (Lynd et al. 2017). Its ability to deconstruct crystalline cellulose is mediated by a protein complex called a cellulosome (Bayer et al. 2008), and this system may have applications for deconstruction of other polymers, including plastics (Yan et al. 2021). In many cases, however, native properties need to be improved for industrial application.

Adaptive laboratory evolution (ALE) is a commonly used strategy for improving desired properties of strains, but experiments can take anywhere from weeks to years (Sandberg et al. 2019). Increasing the mutation rate of a strain can reduce the duration of ALE experiments. Mutations in DNA polymerase III are known to affect the mutation rate of bacteria. DNA polymerase III is responsible for replication of the bacterial genome. It comes in two major forms, DnaE and PolC. The widely-studied *Escherichia coli* only has the DnaE-type enzyme, and many groups have found mutator mutations in this gene (Vandewiele et al. 2002; Strauss et al. 2000; Maki, Mo, and Sekiguchi 1991; Mo, Maki, and Sekiguchi 1991; Ruiz-Rubio and Bridges 1987; Konrad 1978; Sevastopoulos and Glaser 1977). Typically, mutations that affect the fidelity of the DNA Polymerase III holoenzyme are found in the polymerase domain of DnaE or the separate epsilon proofreading subunit.

The PolC-type enzyme is found primarily in gram positive bacteria with low GC content, and has received much less attention (Timinskas et al. 2014). Organisms with PolC typically do not have a separate epsilon proofreading subunit, and instead rely on proofreading activity of the PHP domain within the PolC protein. Several hypermutator mutations in the *polC* gene in Bacillus subtilis have been identified (Barnes et al. 1992) (Gass and Cozzarelli 1973) (Paschalis et al. 2017). A previous ALE experiment identified a *polC* mutation among many mutations in a strain of *C. thermocellum* with a hypermutator phenotype (Holwerda et al. 2020), however the causality of the *polC* mutation was not verified, and the mutation rate was not determined.

In ALE experiments, identifying mutations is only the first step in strain improvement. Mutations have to be subsequently re-introduced so that their effect can be characterized. Previously, it has been difficult to reintroduce point mutations into *C. thermocellum*. Most examples required the deletion of the wild type gene followed by re-introduction of the mutant gene (Zheng et al. 2015; Lo et al. 2015). This process is very time-consuming (~2 months per mutation), and requires that deletion of the target gene is not toxic. Recently, we developed a new CRISPR-based system for introducing point mutations in *C. thermocellum*, based on either the native Type I or heterologous Type II CRISPR systems (Walker et al. 2020).

In this work, we use the newly-developed CRISPR tools to characterize the effect of a single nucleotide mutation in *polC* on the mutation rate of *C. thermocellum*. The ability to both rapidly create diversity with controllable mutator phenotypes, and characterize the resulting mutations with CRISPR tools dramatically improves our ability to perform ALE on *C. thermocellum*.

## Methods

### Plasmid construction

Plasmid construction was performed by isothermal DNA assembly (Gibson 2011), using NEBuilder HiFi DNA Assembly Cloning Kit from NEB. Synthetic DNA was purchased from IDT as gBlocks (Integrated DNA Technologies, Coralville, IA).

### Growth conditions

Routine cultivation was performed using either CTFUD rich medium or MTC-5 chemically defined medium (Olson and Lynd 2012). For solid medium, agar was used at a concentration of 0.8%. Selection for the *cat* marker was performed with 6 ug/ml thiamphenicol. Stock solutions of thiamphenicol were prepared in DMSO at 1000x concentration of 6mg/ml (Sigma, part number:T0251-5G). Selection for the *neo* marker was performed with 150 ug/ml neomycin, stock solution 50 mg/ml in water (Gibco, part number: 21810-031). Selection for *pyrF* mutations was performed with 5-fluoroorotic acid (5-FOA, Zymo Research, part number F9001-5) at a final concentration of 0.5 mg/ml. A 200x 5-FOA stock solution was prepared fresh daily by dissolving 100 mg 5-FOA in 1 ml DMSO. When grown on defined medium, strains with *pyrF* mutations were supplemented with 40 ug/ml uracil (Tripathi et al. 2010), a stock solution of 40 mg/ml uracil was prepared in 1N NaOH (Sigma, part number: U7050-5g).

### Two-step CRISPR approach for introducing mutations

A two-step CRISPR Type I-B approach was used to introduce the *polC* mutation, based on our previously described approach (Walker et al. 2020) with a few modifications. In the two-step approach, first the homology arm plasmid is introduced (primary transformation) and cells are grown to allow homologous recombination to occur. Then a secondary transformation is performed to introduce a killing plasmid to eliminate cells that have not undergone homologous recombination.

#### Primary transformation with homology template

Cultures of parent strain LL1586 (native Type I-B CRISPR system upregulated by insertion of Tsac_0068 promoter) were grown to mid log phase in 50 ml of CTFUD rich medium at 55°C. The culture was centrifuged, rinsed twice with H_2_O, resuspended in H_2_O, transformed using electroporation with plasmid pLT237 containing the *polC* C669Y homology repair template. Transformation was performed using electroporation with a square pulse. The amplitude was 1500 V. A single pulse with a duration of 1.5 ms was applied to a 1 mm cuvette. Cells were removed from the electroporation cuvette and allowed to recover for 16 hours in 2 mL CTFUD at 50°C overnight (Olson and Lynd 2012). The sample was plated on CTFUD/TM (CTFUD medium with 6 ug/ml thiamphenicol) and recovered for 18-24 hours at 55°C. After 5 days, 10 colonies were picked and pooled into 1 mL CTFUD/TM, and incubated overnight at 55°C. The pooled colonies were subcultured twice to provide an opportunity for the homologous recombination events that are selected for in the secondary transformation (Walker et al. 2020). A 1:20 dilution (50 ul into 1 ml) of the pooled colony culture was prepared in CTFUD/TM medium and grown at 55°C overnight for the first transfer. This was repeated for a second transfer. PCR using primers (TY 113+/114+) was used to confirm that plasmid pLT237 was present in the primary culture of 10 pooled colonies, the first transfer culture, and the second transfer culture. All three cultures were all stored at −80°C.

#### Secondary Transformation of LL1586/pLT237 cells with plasmids containing spacers targeting the wildtype *polC* gene

The first and second transfer cultures generated after the primary transformation were pooled and 80 uL of the mixture was inoculated into 10 mL of CTFUD and was grown overnight with and without thiamphenicol (6 ug/mL) to determine the effect of growing the secondary transformation with and without antibiotic selection (note: transformation worked better with added antibiotic at this step, see the results section). These cultures were diluted 1:10 and grown (+/− thiamphenicol) to mid-log phase (Abs_600_ between 0.5 and 0.8), harvested, and transformed as described above. The harvested cells were transformed both individually with plasmids pDGO186N-KS1, pDGO186N-KS2, pDGO186N-KS3, as well as with a pool of the three plasmids, and with a no-DNA control. The plasmids contained a killing spacer (KS1, KS2, or KS3) as well as a neomycin (*neo*) selection marker. After recovery, a portion of the cells were plated on CTFUD/NEO (150 ug/mL neomycin) and incubated at 55°C. Colonies usually appeared after 3 days, and were picked 1-2 days later, resuspended in 0.5 mL of CTFUD/NEO and grown overnight at 55°C. After neomycin selection, colonies from secondary transformations were screened for the target mutation using qPCR.

#### HRM qPCR technique for screening point mutations

The HRM (high-resolution melt analysis) qPCR reaction used a 100 bp amplicon and 25 bp forward and reverse primers that flanked the mutation site (Table 2). For the reaction mixture, 2 uL of bacterial culture was mixed with 10 uL 2X Sso Fast Evagreen qPCR mixture (BioRad USA). The primers were mixed into an equal forward/reverse (F/R) primer mixture and serially diluted to 5 uM. 2 uL of the F/R primer mixture was added along with 6 uL deionized water. The mixture was set to PCR conditions of 95°C/5 min 40X, 95°C/15 s, 55°C/1 min. The range of the melt curve was set to 65°C-95°C at a rate of 0.2°C/10 s. The uAnalyze v2 software (Dwight, Palais, and Wittwer 2012) (Atmadjaja et al. 2019) was used to analyze the raw fluorescence data by making normalized, derivative, and difference plots.

**Table 1.** Strain and plasmids used in this work.

| Strain/Plasmid | Description | Accession Number<sup>a</sup> | Reference |
| --- | --- | --- | --- |
| pLT237 | Replicating plasmid with homology region for introducing PolC<sup>C669Y</sup> mutation. Confers thiamphenicol resistance. | Addgene 174300 | This work |
| pDGO186N-KS1 | Replicating plasmid with sgRNA cassette containing a killing spacer #1 targeting the wild type *polC* gene. | Addgene 174301 | This work |
| pDGO186N-KS2 | Same as above, with killing spacer #2 | Addgene 174302 | This work |
| pDGO186N-KS3 | Same as above, with killing spacer #3 | Addgene 174303 | This work |
| LL1004 | *C. thermocellum* DSMZ 1313 wild type strain; WT *polC*, WT *pyrF* | NCBI reference sequence NC_017304.1 | DSMZ culture collection |
| LL1005 | LL1004 with *pyrF* deletion | not available | (Tripathi et al. 2010) |
| LL1299 | LL1004 with *hpt* and *reIII* deletions | SRX2506395 | (Walker et al. 2020) |
| LL1586 | LL1299 with upregulated cas expression using *Tsac_0068* promoter | SRX4823139 | (Walker et al. 2020) |
| LL1699 | Parent strain LL1586; mutant Pol III, WT *pyrF* | SRX8904888 | This work |
| LL1700 | Parent strain LL1586; mutant Pol III, WT *pyrF* | SRX8904887 | This work |
| LL1738 | LL1586, failed CRISPR mutagenesis at *polC* locus | SRX9642234 | This work |
| LL1740 | LL1586, failed CRISPR mutagenesis at *polC* locus | SRX9642235 | This work |
| LL1741 | LL1586, failed CRISPR mutagenesis at *polC* locus | SRX9642649 | This work |
| LL1742 | LL1586, successful CRISPR mutagenesis, PolC<sup>C669Y</sup> mutation introduced | SRX9642825 | This work |
| LL1743 | LL1586, successful CRISPR mutagenesis, PolC<sup>C669Y</sup> mutation introduced | SRX9642826 | This work |
| LL1744 | LL1586, successful CRISPR mutagenesis, PolC<sup>C669Y</sup> mutation introduced | SRX9642781 | This work |
| LL1745 | LL1586, successful CRISPR mutagenesis, PolC<sup>C669Y</sup> mutation introduced | SRX9642780 | This work |
| LL1746 | LL1586, successful CRISPR mutagenesis, PolC<sup>C669Y</sup> mutation introduced | SRX9642779 | This work |
| LL1762 | LL1742 serial transfer #11 population | SAMN20331610 | This work |
| LL1763 | LL1742 serial transfer #22 population | SAMN20331611 | This work |
| LL1764 | LL1742 serial transfer #33 population | SAMN20331612 | This work |
| LL1765 | LL1745 serial transfer #11 population | SAMN20331613 | This work |
| LL1766 | LL1745 serial transfer #22 population | SAMN20331614 | This work |
| LL1767 | LL1745 serial transfer #33 population | SAMN20331615 | This work |
| LL1768 | LL1738 serial transfer #11 population | SAMN20331616 | This work |
| LL1769 | LL1738 serial transfer #22 population | SAMN20331617 | This work |
| LL1770 | LL1738 serial transfer #33 population | SAMN20331618 | This work |
| LL1793 | LL1738 serial transfer #1 single colony, 50 generations from original | SAMN20331619 | This work |
|  | CRISPR mutant cell |  |  |
| LL1794 | LL1738 serial transfer #11 single colony, 128 generations from original CRISPR mutant cell | SAMN20331620 | This work |
| LL1795 | LL1738 serial transfer #22 single colony, 207 generations from original CRISPR mutant cell | SAMN20331621 | This work |
| LL1796 | LL1738 serial transfer #33 single colony, 286 generations from original CRISPR mutant cell | SAMN20331622 | This work |
| LL1797 | LL1742 serial transfer #1 single colony, 50 generations from original *polC* mutant cell | SAMN20331623 | This work |
| LL1798 | LL1742 serial transfer #11 single colony, 128 generations from original *polC* mutant | SAMN20331624 | This work |
| LL1799 | LL1742 serial transfer #22 single colony, 207 generations from original *polC* mutant cell | SAMN20331625 | This work |
| LL1800 | LL1742 serial transfer #33 single colony, 286 generations from original *polC* mutant cell | SAMN20331626 | This work |
| LL1801 | LL1745 serial transfer #1 single colony, 50 generations from original *polC* mutant cell | SAMN20331627 | This work |
| LL1802 | LL1745 serial transfer #11 single colony, 128 generations from original *polC* mutant cell | SAMN20331628 | This work |
| LL1803 | LL1745 serial transfer #22 single colony, 207 generations from original *polC* mutant cell | SAMN20331629 | This work |
| LL1804 | LL1745 serial transfer #33 single colony, 286 generations from original *polC* mutant cell | SAMN20331630 | This work |

<sup>a</sup> Accession numbers that begin with “SRX” represent samples with WGS performed at JGI, numbers that begin with “SAMN” represent samples with WGS performed at Dartmouth. Both sets are available from the NCBI SRA database.

**Table 2.**
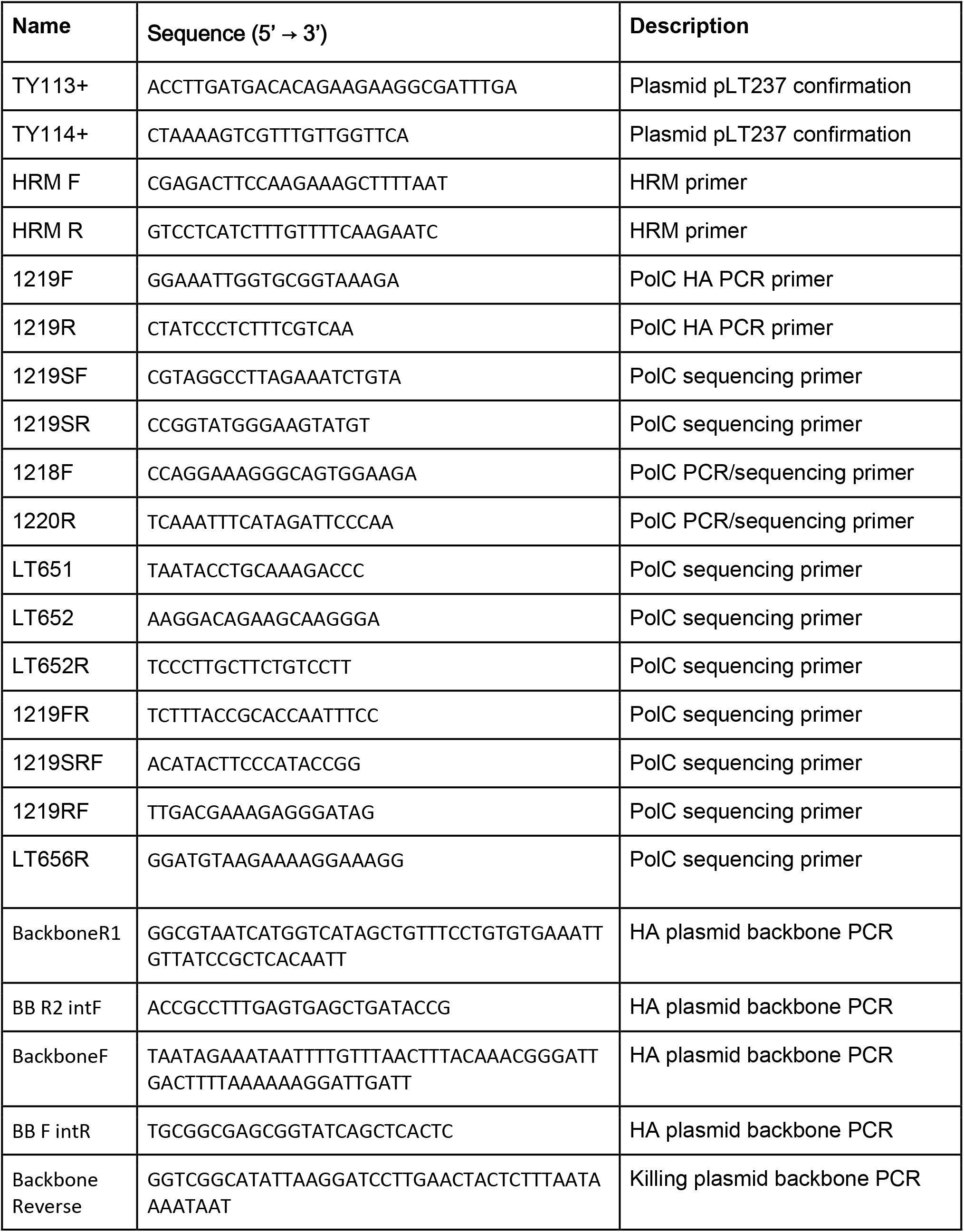
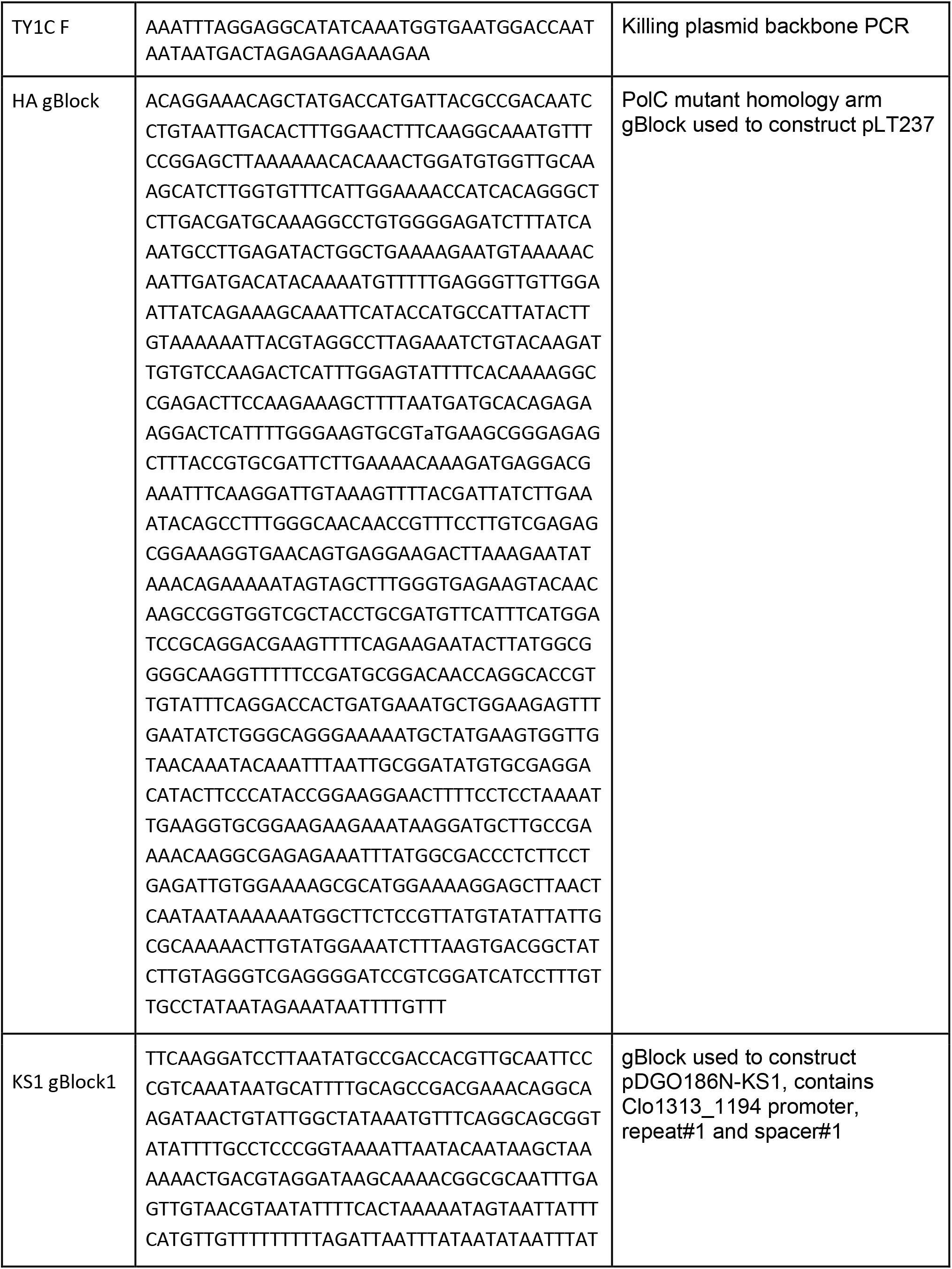
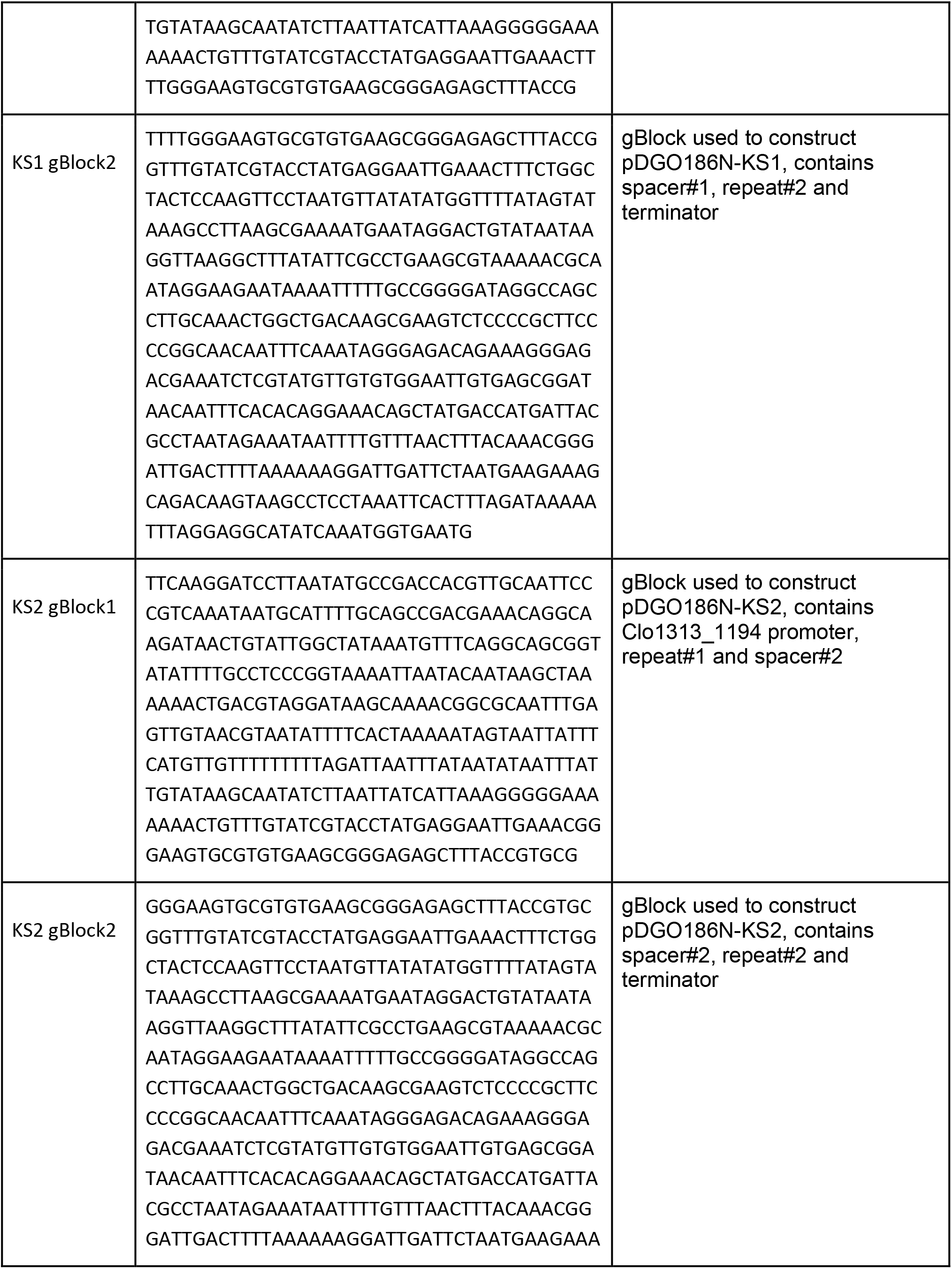
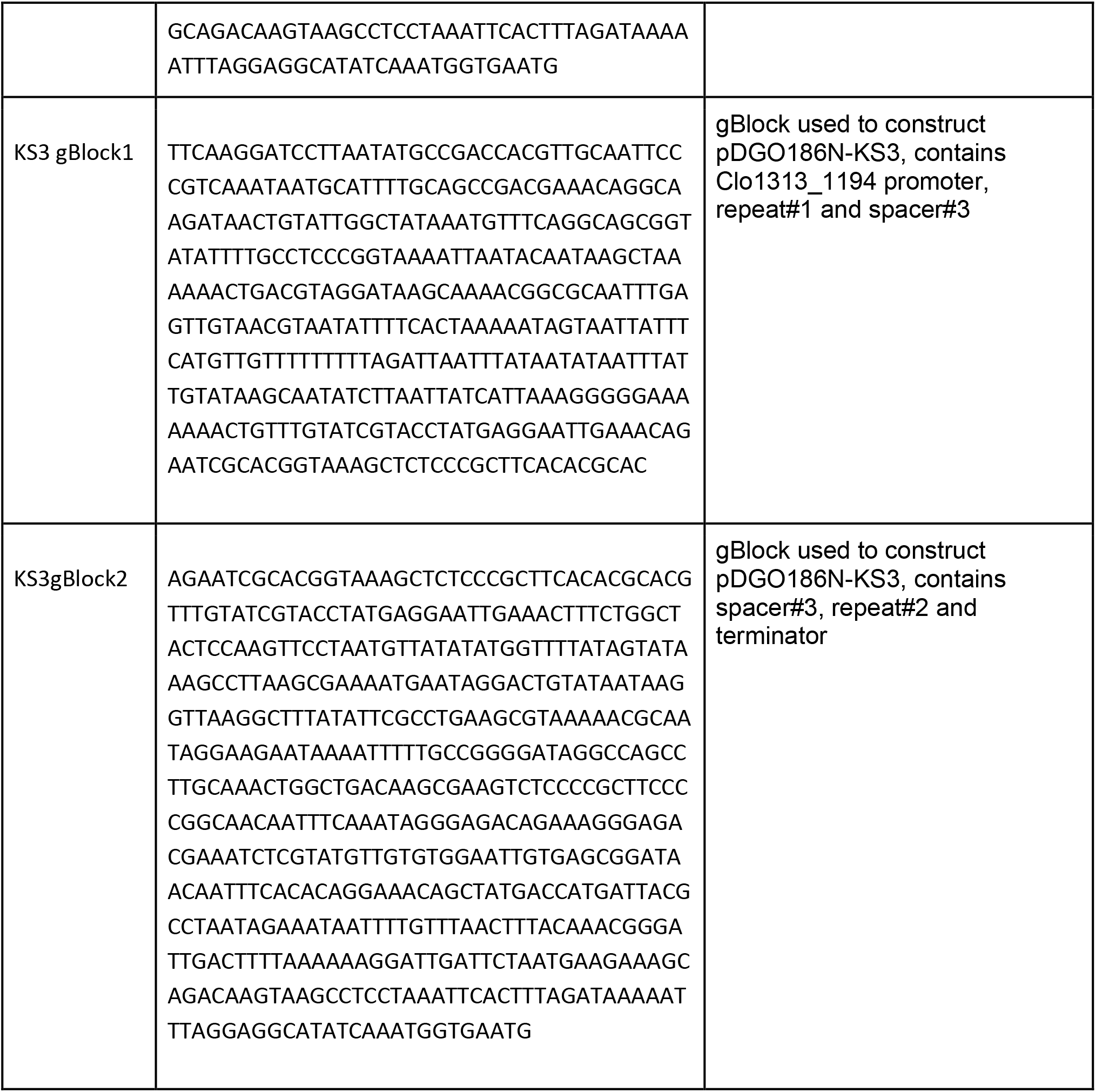
Primers and synthetic DNA (gBlocks) used in this work.

### 5-FOA resistance test

For the 5-FOA resistance test, 5 to 50 ul of a freezer stock, was inoculated into 1.5 ml media, grown for 6-8 hours to mid-log phase (to ensure a rapidly-dividing culture, which is optimal for 5-FOA selection), and 100 ul was plated at various dilutions with and without 5-FOA to ensure that there were between 10-200 colonies on each plate. Usually the 10^−4^, 10^−5^ and 10^−6^ dilutions had a countable number of colonies for the 5-FOA plates. The plates were incubated at 55°C for 4-6 days, and colonies were counted to determine the fraction of cells exhibiting 5-FOA resistance.

### Mutation accumulation experiment

To determine the mutation rate by mutation accumulation, cultures were serially transferred. In bacterial cells, single colony isolation events usually correspond to about 23-25 generations (Trindade, Perfeito, and Gordo 2010; Kibota and Lynch 1996). For *C. thermocellum*, based on cell volume (0.4 um diameter by 2 um long (Sato et al. 1993)), and colony volume (lenticular shape approximately 1 mm in diameter by 0.4 mm in height), a colony should contain about 5e8 cells, corresponding to 29 generations. Approximately another 18 generations (1:100 dilution from initial culture of colony pick and 1:2500 dilution subculture for freezer stock preparation) occurred between the initial colony isolation following the introduction of the *polC* mutation and the preparation of the freezer stock. This does not affect the mutations observed in the starting strains (L1699, LL1700, LL1738, LL1740-LL1746), but needs to be considered for the subsequent single colony isolations. Each serial transfer consisted of a 1:100 dilution (~6.6 generations). After 10 1:100 serial transfers, the 11th transfer was a 1:5000 dilution (~12.3 generations) to provide extra volume for preparing freezer stocks and gDNA for WGS. This was repeated three times. The 11th, 22nd, and 33rd transfers were stored as populations in the freezer, and the presence of the *polC* mutation was confirmed by HRM qPCR.

Single colonies were isolated from each population to create a population bottleneck to fix mutations. This involved another 1:100 subculture (~6.6 generations), followed by plating on solid medium. A single colony was picked, grown up, and prepared for whole genome sequencing (WGS).

### Whole genome resequencing (WGS) at Dartmouth

Genomic DNA was prepared using the Omega E.Z.N.A. kit following the manufacturer’s protocol (Omega Bio-Tek, GA, USA). 500 ng of DNA was used for WGS library preparation using the NEBNext Ultra II FS DNA Library Prep Kit for Illumina (New England Biolabs, MA, USA). Fractionated, adapter ligated DNA fragments went through 5 rounds of PCR amplification and purification. The resulting WGS library was sequenced at the Genomics and Molecular Biology Shared Resource (GMBSR) at Dartmouth. Libraries were diluted to 4 nM, pooled and loaded at 1.8 pM onto a NextSeq500 Mid Output flow cell, targeting 130 million 2×150 bp reads/sample. Base-calling was performed on-instrument using RTA2 and bcls converted to fastq files using bcl2fastq2 v2.20.0.422.

### Whole genome resequencing (WGS) at JGI

Genomic DNA was submitted to the Joint Genome Institute (JGI) for sequencing with an Illumina MiSeq instrument. Paired-end reads were generated, with an average read length of 150 bp and paired distance of 500 bp. Unamplified libraries were generated using a modified version of Illumina’s standard protocol. 100 ng of DNA was sheared to 500 bp using a focused ultrasonicator (Covaris). The sheared DNA fragments were size selected using SPRI beads (Beckman Coulter). The selected fragments were then end repaired, A-tailed and ligated to Illumina compatible adapters (IDT, Inc) using KAPA Illumina library creation kit (KAPA biosystems). Libraries were quantified using KAPA Biosystem’s next-generation sequencing library qPCR kit and run on a Roche LightCycler 480 real-time PCR instrument. The quantified libraries were then multiplexed into pools for sequencing. The pools were loaded and sequenced on the Illumina MiSeq sequencing platform utilizing a MiSeq Reagent Kit v2 (300 cycle) following a 2 × 150 indexed run recipe.

### WGS data analysis

Read data was analyzed with the CLC Genomic Workbench version 12 (Qiagen Inc., Hilden, Germany). First, reads were trimmed using a quality limit of 0.05 and ambiguity limit of 2. Then 2.5M reads were randomly selected (to avoid errors due to differences in the total number of reads). Reads were mapped to the reference genome (NC_017992). Mapping was improved by two rounds of local realignment. The CLC Basic Variant Detection algorithm was used to determine small mutations (single and multiple nucleotide polymorphisms, short insertions and short deletions). Variants occurring in less than 35 % of the reads or fewer than 4 reads were filtered out. The fraction of the reads containing the mutation is presented in XXX . To determine larger mutations, the CLC InDel and Structural Variant algorithm was run. This tool analyzes unaligned ends of reads and annotates regions where a structural variation may have occurred, which are called breakpoints. Since the read length averaged 150 bp and the minimum mapping fraction was 0.5, a breakpoint can have up to 75 bp of sequence data. The resulting break-points were filtered to eliminate those with fewer than ten reads or less than 20 % “not perfectly matched.” The breakpoint sequence was searched with the Basic Local Alignment Search Tool (BLAST) algorithm (Altschul et al. 1990) for similarity to known sequences. Pairs of matching left and right breakpoints were considered evidence for structural variations such as transposon insertions and gene deletions. The fraction of the reads supporting the mutation (left and right breakpoints averaged) is presented in Supplementary Table 1. Mutation data from CLC was further processed using custom Python scripts (https://github.com/danolson1/cth-mutation).

### Sanger sequencing

Colony PCR was performed on bacterial cultures and the PCR product purified using the DNA Clean and Concentrator kit (Zymo). Purified PCR products were sequenced at Genewiz (USA).

### Quantification of mutation rate

The mutation rate was determined based on synonymous mutations, which are generally assumed to have a low effect on fitness (M Kimura 1968). *C. thermocellum* has a genome size of 3,561,619 bp (NC_017304.1) of which 83% consists of coding regions. For each codon, the number of synonymous single-substitution events were counted, and multiplied by the codon frequency (Kazusa Codon Usage Database) (Nakamura and Tabata 1997), to reveal that 21.4% of all nucleotide positions in *C. thermocellum* coding sequences allow a synonymous mutation. This results in an effective genome size of 632,024 bp. The mutation rate (□, mutations per base pair per generation) is calculated with the following equation:

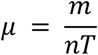

Where m is the number of observed mutations, n is the number of sites analyzed on the genome, and T is the number of generations (Kucukyildirim et al. 2020; Lynch et al. 2016).

## Results and Discussion

### CRISPR system successfully introduces point mutations

Initially, we attempted to re-introduce the C669Y mutation into the *polC* gene using standard homologous recombination techniques (Olson and Lynd 2012). This required cloning the entire *polC* gene to ensure that a functional copy was present at each stage of the chromosomal modification. Despite repeated attempts, we were unable to construct the deletion vector due to apparent toxicity in *E. coli*.

We thus pursued an alternative approach, using a recently-developed chromosomal modification technique that co-opts the native Type I CRISPR system in *C. thermocellum* (Walker et al. 2020). This system involves two transformation events (Fig. 1). The first transformation introduces the homology repair template, which introduces the desired point mutation, as well as several silent mutations to prevent spacer recognition. The second transformation introduces the killing spacer module, which targets chromosomes with the wild type *polC* sequence, but not ones modified by the homology repair template. Colonies were screened for the presence of the *polC* mutation using an HRM PCR assay (Supplementary Figure 1), which identifies mutations by a decrease in the melting temperature of mutant PCR amplicons. After transformation with the killing spacer plasmid, colonies were picked, and the *polC* region was analyzed by HRM qPCR. A total of 309 colonies were picked for HRM qPCR screening and 17 mutant candidates were identified (Table 3), Sanger DNA sequencing confirmed the presence of the *polC* mutation in all 17 candidates. Performing the secondary transformation with cells grown in the presence of thiamphenicol resulted in 5-fold higher transformation efficiency. Mutations were subsequently confirmed by Sanger sequencing and whole-genome sequencing, and 15 of 159 colonies (9.4%) had the correct mutant genotype. In most strains with the *polC* mutation, both silent mutations were also present. One strain (LL1746) was missing one of the silent mutations, indicating that both silent mutations are not necessary to prevent CRISPR targeting.

**Figure 1.**
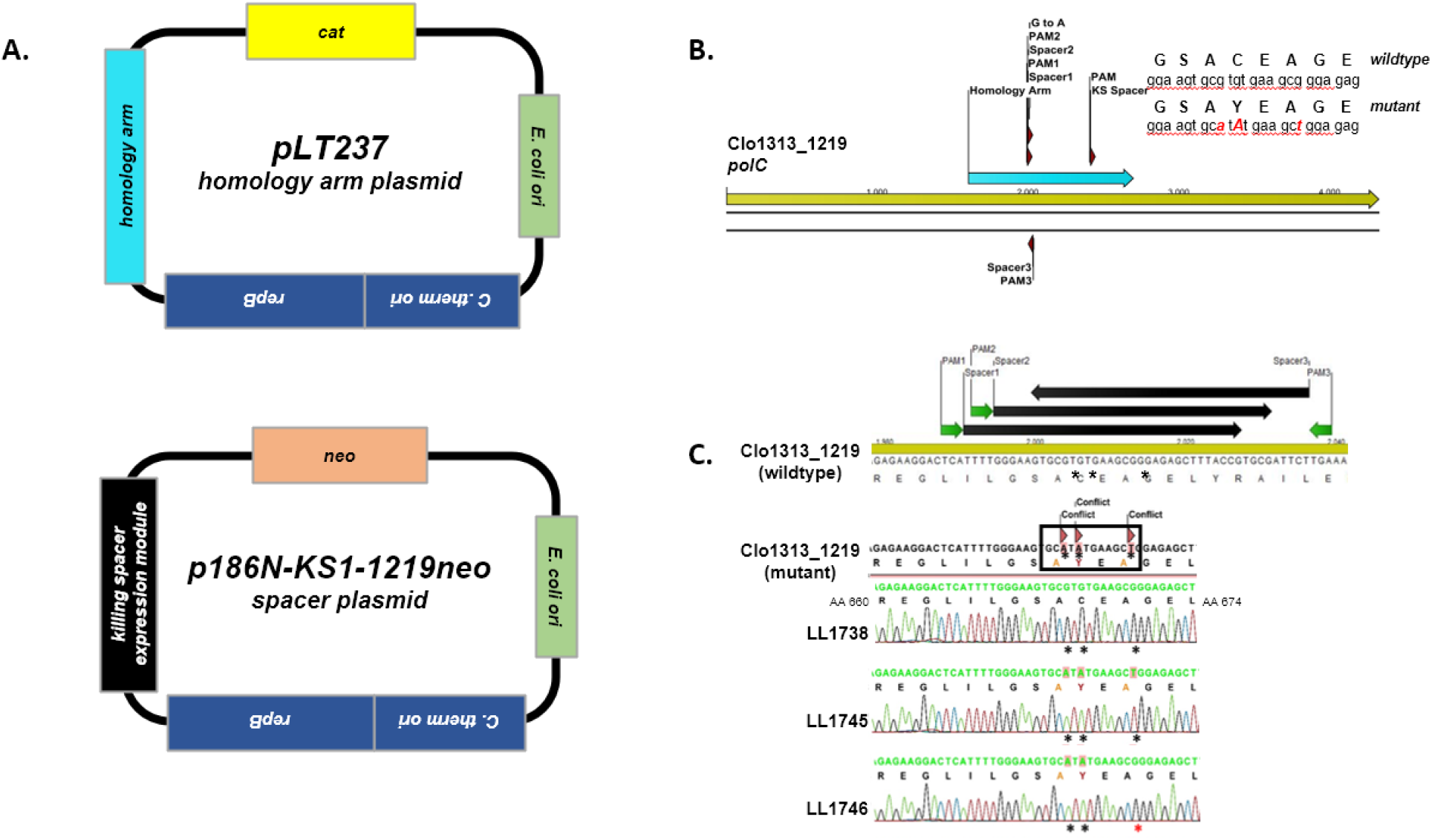
**A) 2-step CRISPR system for introducing mutations in *C. thermocellum***. Step 1 is transformation with a homology arm plasmid, and step 2 is transformation with a killing spacer cassette plasmid. Each plasmid contains a *C. thermocellum* origin (including the *repB* replication protein), an *E. coli* origin, and either a chloramphenicol (*cat*) or a neomycin (*neo*) selection marker. **B) The *polC* region targeted for mutations.** The homology arm is shown at the start of the target sequence. The spacer cassette is composed of three spacers oriented in the forward (KS1 or KS2) or reverse (KS3) directions. Spacer regions were chosen to be immediately downstream of a TTN or TCD PAM sequence. The three target mutations are also shown: two silent mutations (to disrupt CRISPR targeting) and the targeted point mutation to create a cysteine to tyrosine amino acid change at position 669. **C) Confirmation of mutagenesis.** The *polC* mutation targets were analyzed by Sanger sequencing. Strain LL1738 is an example of failed mutagenesis, and displays a wild-type sequence. Strain LL1745 is an example of successful mutagenesis, exhibiting all three targeted mutations. In strain LL1746, the *polC* mutation was introduced successfully with only one of the two silent mutations.

**Table 3.**
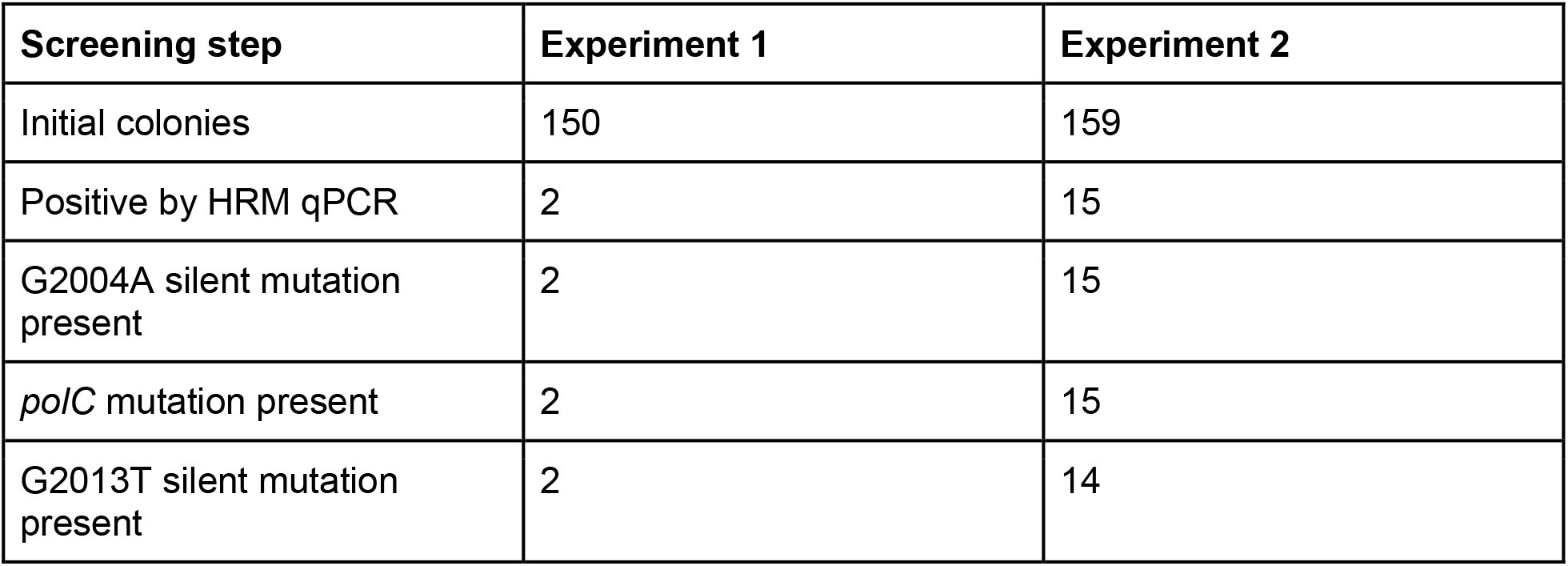
HRM screening results

The genome editing work was performed at the same time as some of the experiments in our previous publication on CRISPR-based editing in *C. thermocellum* (Walker et al. 2020), and thus does not include several improvements described in that work, such as the incorporation of thermostable recombinases (*exo* and *beta* genes from *Acidithiobacillus caldus*).

To eliminate CRISPR-mediated restriction, we opted to make two silent mutations in the spacer region, rather than disrupt the PAM sequence (Fig. 1). A benefit of this approach is that we can use a single homology arm plasmid (first transformation) with several killing spacer plasmids (second transformation), however the editing efficiency is slightly lower than what we previously reported (10% vs 40%) (Walker et al. 2020).

The effect of the PolC^C669Y^ mutation was determined with a 5-FOA resistance assay (Fig. 2). In rich media, the *de novo* pyrimidine biosynthesis pathway is non-essential, and it is thus a neutral site where mutations can accumulate. Mutations that inactivate this pathway create a 5-FOA resistant (Foa^r^) phenotype that can readily be detected, and this assay is frequently used as a measure of mutation rates (Boeke, La Croute, and Fink 1984; Grogan and Gunsalus 1993; Kondo, Yamagishi, and Oshima 1991; Jacobs and Grogan 1997). In this experiment, we observed a natural abundance of the Foa^r^ phenotype at about 2% of WT cells, similar to what we have reported previously for *C. thermocellum* (Tripathi et al. 2010). Disruption of *pyrF* largely eliminates sensitivity to 5-FOA, with more than 70% of colonies exhibiting the Foa^r^ phenotype (the reason this number is not 100% is likely due to a decrease in plating efficiency in the selective condition). Most of the strains harboring the PolC^C669Y^ mutation also show an increase in the Foa^r^ phenotype, however this was not universally observed. Strain LL1700 has the PolC^C669Y^ mutation, but is Foa^s^, and strain LL1740 does not have the PolC^C669Y^ mutation, but is Foa^r^.

**Figure 2.**
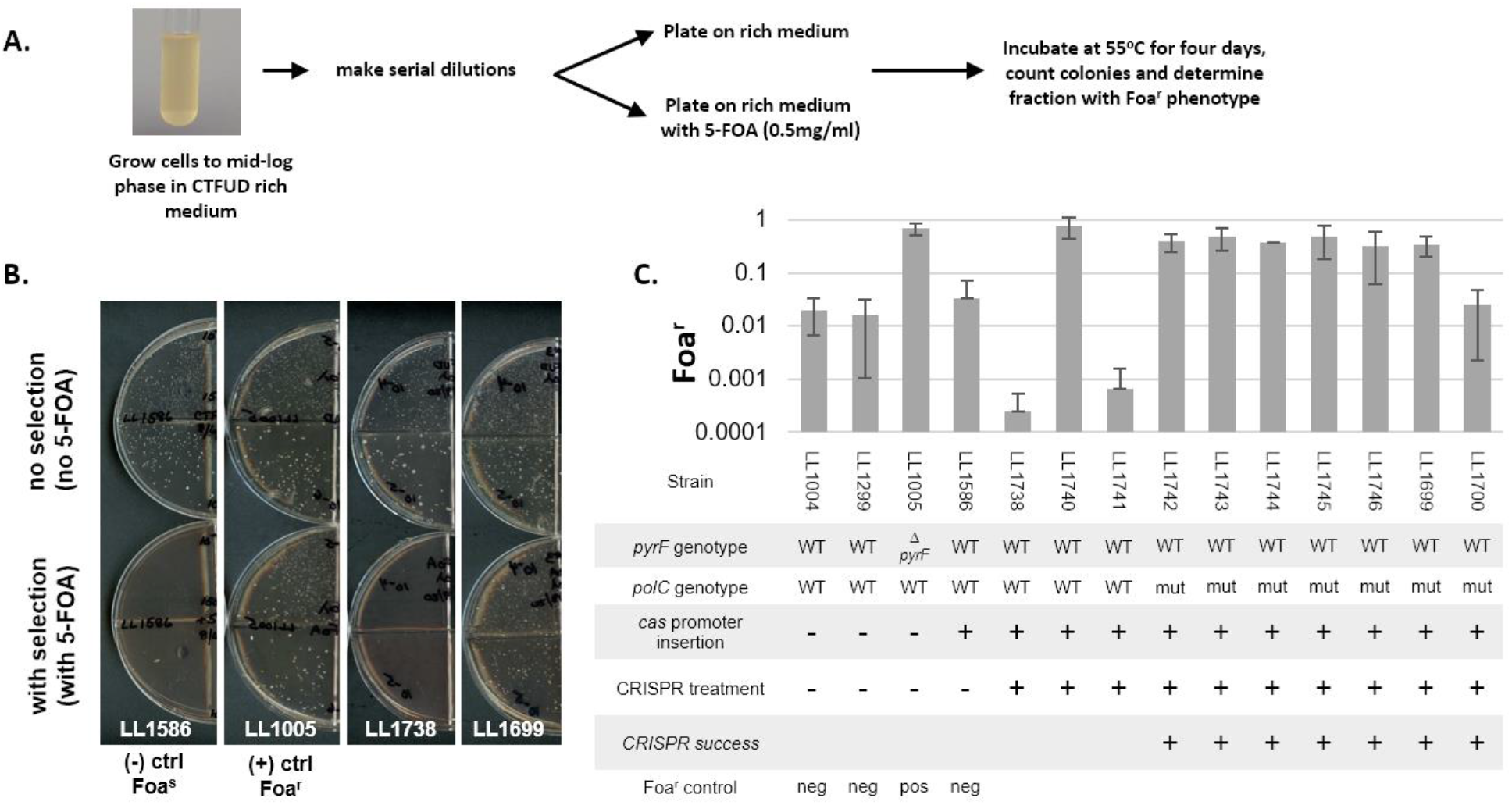
5-FOA resistance mutation rate assay. **A)** Schematic diagram of cell growth and selective plating. **B)** Dilution plates in the selective (with 5-FOA) and permissive (no 5-FOA) conditions. The parent strain (LL1586) is WT at the *pyrF* locus and Foa^s^. Strain LL1005 has a targeted *pyrF* deletion and is Foa^r^. Strain LL1738 exhibits the Foa^s^ phenotype. Strain LL1699 exhibits the Foa^r^ phenotype. **C)** Foa^r^ phenotype for wild-type and mutant *polC* strains. The *pyrF* genotype, *polC* genotype, *cas* promoter insertion, and success of CRISPR treatment were determined by Sanger sequencing.

In order to calculate the mutation rate, the target size needs to be known. We expected to observe mutations primarily in the *pyrF* gene, however Sanger sequencing did not reveal the expected mutations at that locus. Initially we plated on CTFUD with 5-FOA, picked 12 LL1299 (WT) colonies and sequenced the *pyrF* gene, but did not find any mutations. We sequenced the *pyrF* gene from 40 LL1699 (*polC* mutant) colonies picked from 5-FOA plates, and only found one mutation (a single nucleotide insertion resulting in a frame shift). Since *pyrF* mutants exhibit a growth defect that can be complemented with added uracil (Tripathi et al. 2010), we repeated the LL1299 plating experiment on CTFUD with 5-FOA and 40 ug/ml uracil. Of 18 colonies, 5 had mutations (mostly frame-shifts and premature stop codons) at the *pyrF* locus.

Subsequent whole-genome sequencing (Supplementary Table 1) did not identify any mutations associated with *de novo* pyrimidine biosynthesis, and the genetic basis for the Foa^r^ phenotype in these strains remains unknown, which prevents an accurate determination of the mutation rate. Instead, we decided to measure mutation accumulation directly, by whole-genome sequencing.

For this mutation accumulation experiment, three strains were selected that had undergone CRISPR mutagenesis: one where the mutagenesis had failed (strain LL1738, WT at *polC*) to serve as a control, and two where the CRISPR mutagenesis had been successful (strains LL1742 and LL1745). Each strain was serially transferred for several generations (Fig. 3). Every 11 generations, the population was stored in the freezer, and a single colony was isolated. Observing the accumulation of mutations over time allows us to estimate the mutation rate.

**Figure 3.**
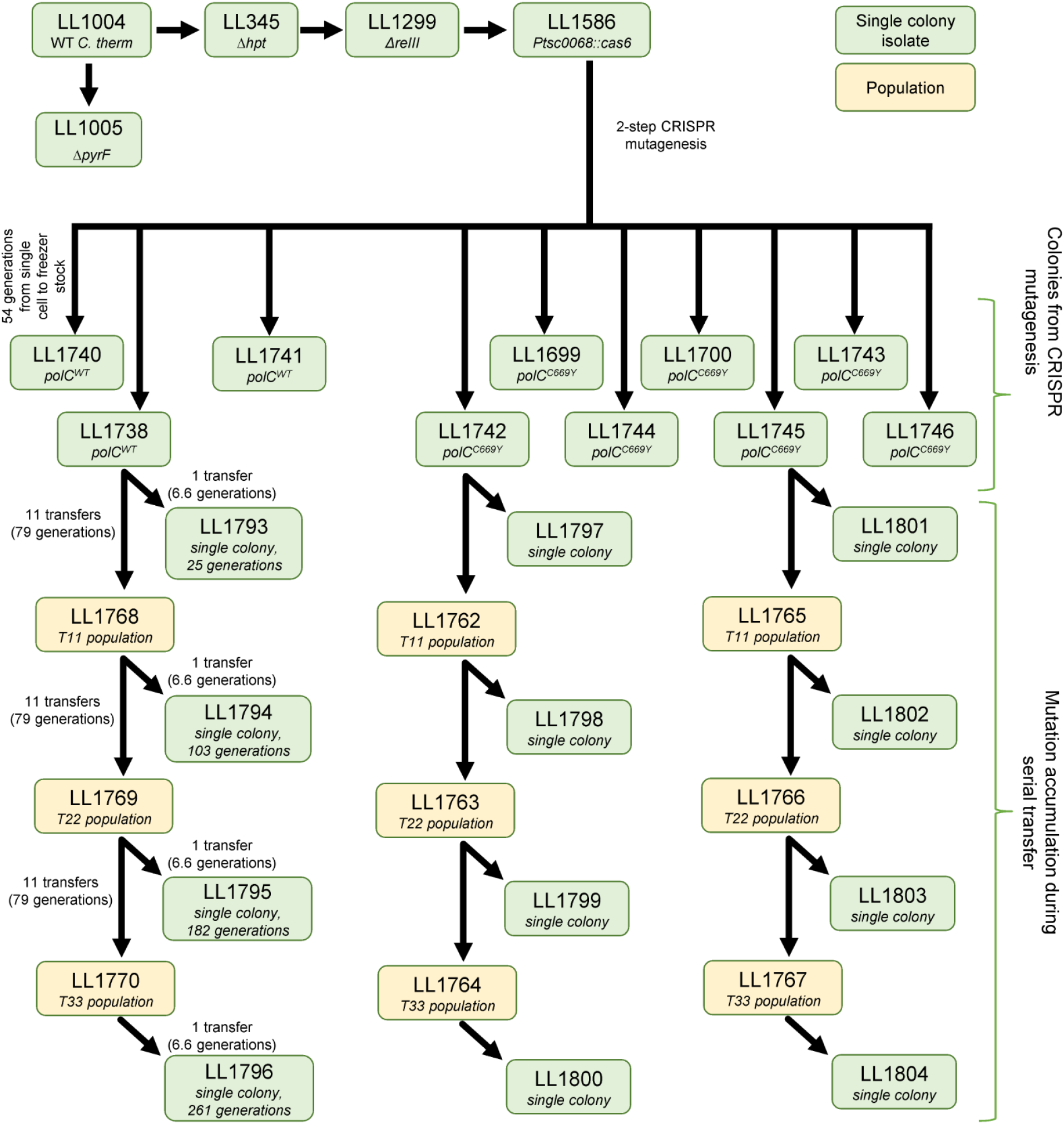
Lineages of the strains described in this work. Strain LL1586 was derived from WT *C. thermocellum* (LL1004) by a series of targeted mutations designed to improve the ability to perform targeted genetic modifications. Two-step CRISPR mutagenesis was performed on strain LL1586 to introduce the PolC^C669Y^ mutation. Eight individual colonies were isolated and sequenced at the *polC* locus.

Whole-genome sequencing indicated a substantial increase in the rate of mutation accumulation for strains with the PolC^C669Y^ mutation (Fig. 4). Several categories of mutations were identified: One category is structural rearrangements, which are mutations that involve large (> 3 nucleotide) insertions, deletions, or replacements. This includes the targeted genetic modifications (deletion of *hpt* in strain LL345 (Argyros et al. 2011), deletion of *reIII* in strain LL1299 (Riley et al. 2019), insertion of a strong constitutive promoter from *Thermoanaerobacterium saccharolyticum* driving the native Type I cas operon in strain LL1568 (Walker et al. 2020). This also includes transposon insertions. *C. thermocellum* has several native transposon elements (Zverlov et al. 2008; Holwerda et al. 2020). Transposons insertions from families IS2, IS10, and IS120 (Siguier et al. 2006) were observed. All of these insertions, except for two in the LL1769 population, were inherited from the LL1586 parent strain. The two in the LL1769 population were not present in either the subsequent serial transfer (LL1770 population) or the single colony isolate from that transfer (strain LL1795), and we therefore suspect it appeared during the preparation for genomic DNA extraction for whole-genome resequencing. Mutations in this category comprise about 1% of the total number of mutations (7 of 775 for the 12 single colony isolates in the mutation accumulation experiment). Since these mutations are not expected to be affected by PolC mutations, they were excluded from subsequent analysis.

**Figure 4.**
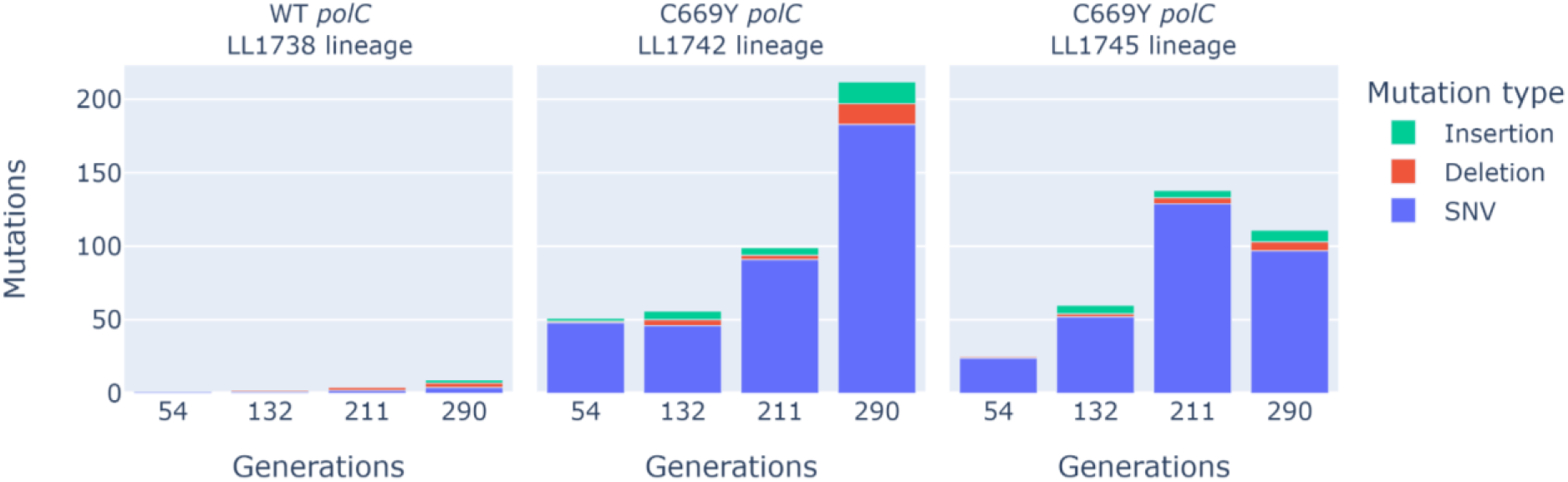
Accumulation of mutations identified by whole-genome resequencing. Each transfer is a 1:100 dilution. For each lineage, mutations present in the parent strain were filtered out. SNV stands for “single nucleotide variation.” The data used to generate this figure is presented in detail in Supplementary Table 1.

Another category is short (1-3 nucleotide) insertions, deletions, and replacements. These comprise the majority (99%) of observed mutations, and the incidence of these mutations was elevated in PolC mutant strains (Fig. 4).

The mutation distribution is not uniform, with an underrepresentation of A:T → T:A and G:C → C:G transversion relative to other types of transitions and transversions, and an overrepresentation of C:G → A:T transversions (Figure 5). This tendency has been observed by others, however a mechanism is not known (Wielgoss et al. 2011; Hershberg and Petrov 2010). Nevertheless, it is important to take this into consideration when designing adaptive laboratory evolution experiments, since this mutational bias affects the likelihood of observing amino acid changes.

**Figure 5.**
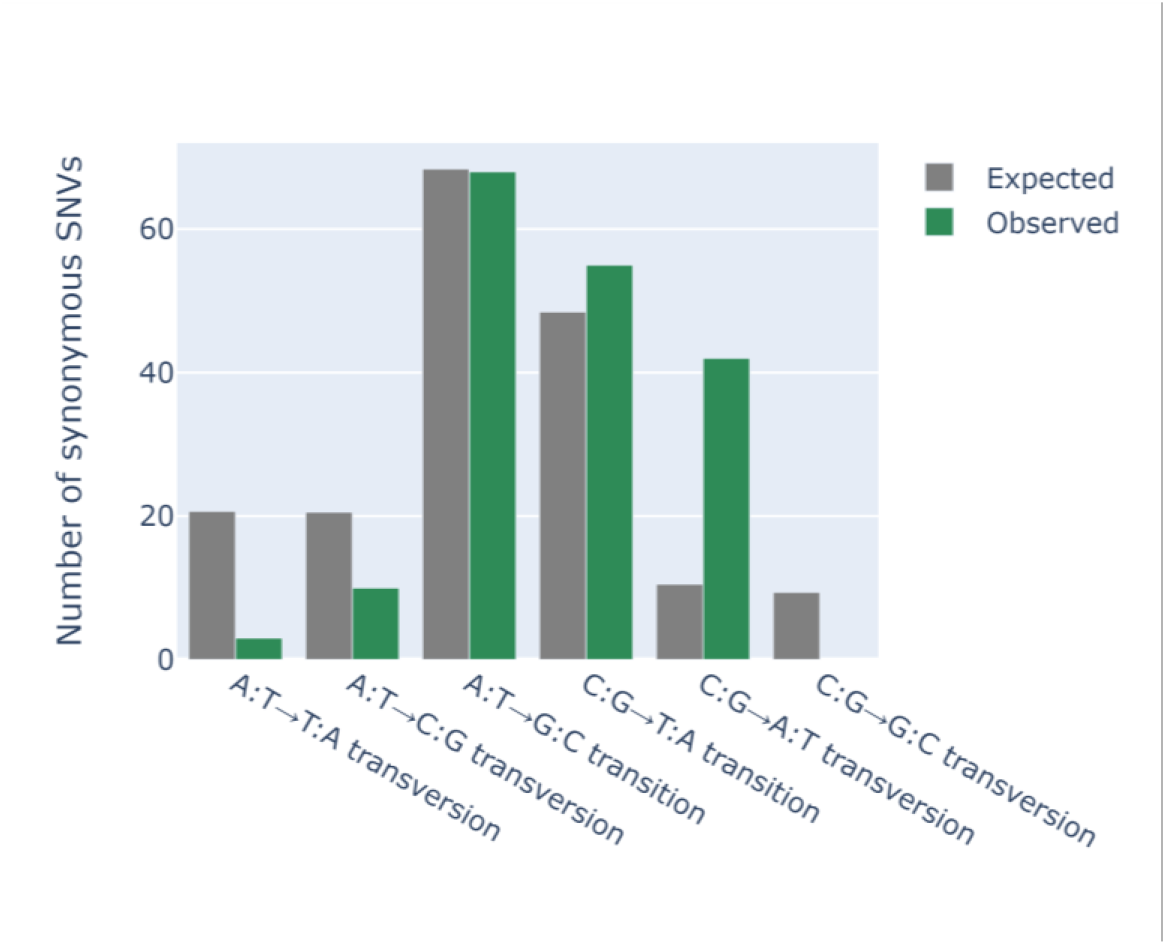
Comparison of mutation types. For this analysis, only synonymous SNV mutations from the *polC* mutant strain lineages (strains LL1797, LL1798, LL1799, LL1800, LL1801, LL1802, LL1803, and LL1804) were considered. The wild type strains did not have any synonymous mutations. Expected mutation frequency was determined based on the codon frequency and the probability that a mutation results in a synonymous mutation.

### Quantification of the mutation rate

The simplest way to calculate the mutation rate is to divide the total number of mutations by the total number of generations. However, some mutations are lethal and thus not observed. To correct for this, we can use synonmyous SNVs, a subset of mutations that are presumed to be neutral (Motoo Kimura 1984). In the *polC* mutant strains, the median mutation rate is 0.10 mutations per generation or 1.6e-7 per nucleotide. It is not possible to determine the mutation rate for the WT strain using synonymous mutations, since even after 290 generations, no synonymous mutations were observed. We can, however, establish an upper bound of about 5.5e-9 per nucleotide (assuming a single mutation appeared after 290 generations). We can also estimate the upper bound considering all four non-synonymous mutations in strain LL1796 (WT PolC, transfer 33). This gives a mutation rate of 3.9e-9 per generation. Many organisms have a mutation rate of about 0.0033 per generation (Drake 1991), which would be 9.2e-10 for an organism with the genome size of *C. thermocellum*. Thus, the *polC* mutation appears to have increased the mutation rate between 30 and 178-fold, and looking at the structure of this enzyme suggests a possible mechanism.

### C669Y mutation may disrupt proofreading activity of *polC*

DNA polymerase III comes in two major forms: DnaE, and PolC. The DnaE types are further divided into three subtypes (DnaE1, DnaE2, and DnaE3). *C. thermocellum* contains both a PolC type (Clo1313_1219) and a DnaE1 type (Clo1313_0994), both of which are expressed (Holwerda et al. 2020). Many organisms with PolC, including *C. thermocellum*, do not have an epsilon proofreading subunit. In these organisms, proofreading is mediated either by the embedded EXO domain or by the PHP domain itself. This domain has been shown to have proofreading activity in *Thermus thermophilus* (Stano, Chen, and McHenry 2006). The C669Y mutation is located in the PHP domain of PolC (Fig. 6). Exonuclease activity depends on coordination with several metal ions via nine highly conserved residues. (Timinskas et al. 2014) The C669Y mutation disrupts one of these residues, which may subsequently disrupt metal ion binding and thus impair proofreading activity.

**Figure 6.**
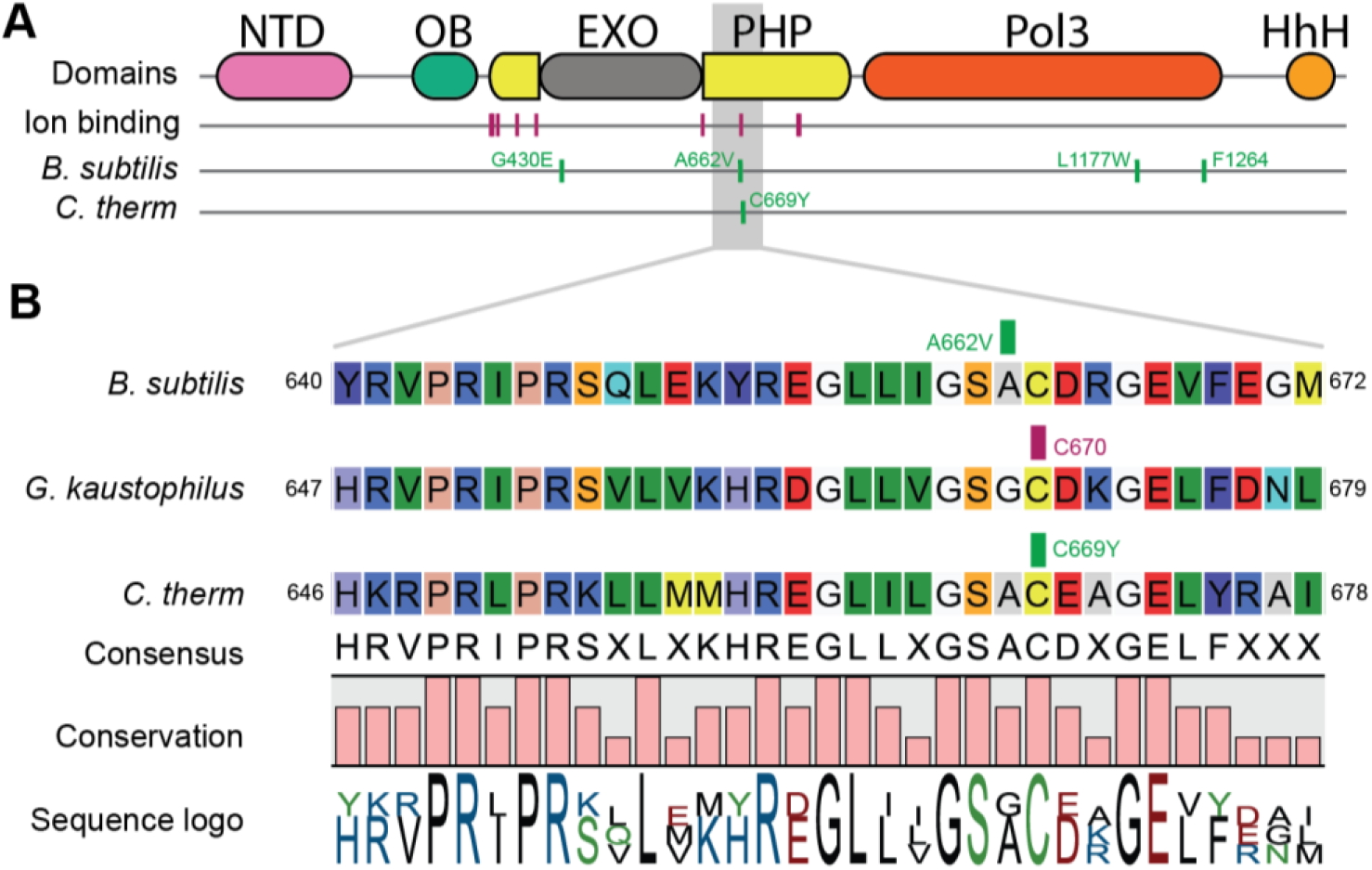
Location of the PolC^C669Y^ mutation. **Panel A** shows the domain structure of the PolC protein. The C669Y mutation is located in the PHP (polymerase and histidinol phosphatase) domain. Other domains include N-terminal domain (NTD), oligonucleotide binding (OB), exonuclease (EXO), polymerase core (Pol3), and tandem helix-hairpin-helix motif (HhH). The PHP domain contains eight highly conserved residues (magenta) that coordinate binding with the metal ions, Mn^2+^ and Zn^2+^. Mutations known to affect polymerase fidelity in *B. subtilis* are also indicated. **Panel B** shows a detailed view of the region surrounding the C669Y mutation. The C669Y mutation is adjacent to an A662Y mutation observed to cause a temperature-sensitive phenotype in *B. subtilis*. The C669 residue in *C. thermocellum* is in the same position as the C670 residue in *G. kaustophilus* (PDB ID: 3F2D) that coordinates the Mn^2+^ residue.

Similar mutations have been observed in *Bacillus* and *E. coli*. In *B. subtilis*, the A662V mutation is adjacent to the metal-binding cysteine. It does not have an effect on the mutation rate, but *does* cause temperature instability. (Barnes et al. 1992) In *E. coli*, the *dnaE74* mutation (G134R) is in a similar location (Vandewiele et al. 2002), however in *E. coli*, the PHP domain does not have any of the conserved metal-binding residues, and is not thought to have catalytic activity. The observed change in mutation rate (1.8-fold increase) in that organism may be due to changes in binding affinity for the epsilon proofreading subunit (*dnaQ*) instead.

### Other mutations that may affect the mutation rate

In addition to the *polC* mutation, several other mutations were identified that might affect the mutation rate. Strain LL1803 (transfer 22 in the LL1745 lineage) has a mutation in the *mutS* gene (Clo1313_1201) responsible for DNA mismatch repair. Mutations in this gene have been shown to increase the mutation rate in other organisms (Luan et al. 2013; Willems et al. 2003). Strain LL1797 (transfer 1 in the LL1742 lineage) has a mutation in the DNA polymerase III delta subunit (Clo1313_1173). The delta subunit is part of the DNA polymerase III holoenzyme, and is responsible for opening the sliding beta clamp protein to accept a DNA strand for replication (Turner et al. 1999). Strain LL1800 (transfer 33 in the LL1742 lineage) has a mutation in the *polA* polymerase (Clo1313_1334). This polymerase is thought to assist in repairing damaged DNA (Hernández-Tamayo et al. 2019). Several strains (LL1801, LL1803, and LL1804) have mutations in the *lexA* DNA binding protein (Clo1313_2881). All three mutations are in different locations in the gene, and two of them likely eliminate activity (one is a frameshift mutation, the other is a stop codon). The *lexA* gene works in tandem with the *recA* gene to induce the SOS response (programmed DNA repair) (Butala, Zgur-Bertok, and Busby 2009). Since *lexA* is a repressor of SOS activity, its inactivation by mutation would be expected to lead to constitutive induction of the SOS response. Although the effect of these mutations has not been tested, at least some of them are likely to be anti-mutator alleles. Hypermutator phenotypes that arise in bacterial populations typically revert to the ancestral mutation rate when maintained in stable conditions (Couce et al. 2017).

## Conclusions

We demonstrate the utility of our recently developed CRISPR/cas system to successfully introduce a PolC^C669Y^ mutation in *C. thermocellum*. This is the first example of the use of a CRISPR system to introduce a *novel* mutation identified by ALE (the Walker et al. 2020 paper demonstrated proof-of-concept by putting a stop codon in the *pyrF* gene). We found the HRM technique to be useful for rapidly screening colonies to identify the successful introduction of point mutations. The single C669Y mutation in PolC protein in *C. thermocellum* is sufficient to increase the mutation rate about 30-fold. This mutation appears to function by interfering with metal ion coordination in the PHP domain responsible for proofreading. The ability to selectively increase the mutation rate in *C. thermocellum* is a useful tool for directed evolution experiments.

## Acknowledgements

We thank Dr. Česlovas Venclovas for useful discussions related to DNA polymerases. Funding was provided by The Center for Bioenergy Innovation, a U.S. Department of Energy Research Center supported by the Office of Biological and Environmental Research in the DOE Office of Science. Whole genome resequencing was performed by the Department of Energy Joint Genome Institute, a DOE Office of Science User Facility, and is supported by the Office of Science of the U.S. Department of Energy under contract number DE-AC02–05CH11231. Additional whole genome resequencing was carried out in the Genomics and Molecular Biology Shared Resource (GMBSR) at Dartmouth which is supported by NCI Cancer Center Support Grant 5P30CA023108

Lee R. Lynd is a cofounder of the Enchi corporation, a start-up company focusing on cellulosic ethanol production using Clostridium thermocellum. There are no other competing interests.

